# Genome-wide analysis of lncRNAs and mRNAs expression during the differentiation of abdominal preadipocytes in chicken

**DOI:** 10.1101/069591

**Authors:** Tao Zhang, Xiangqian Zhang, Kunpeng Han, Genxi Zhang, Jinyu Wang, Kaizhou Xie, Qian Xue

## Abstract

lncRNAs regulate metabolic tissue development and function, including adipogenesis. However, little is known about the function and profile of lncRNAs in preadipocytes differentiation of chicken. Here, we identified lncRNAs in preadipocytes of different differentiation stages by RNA-sequencing using Jinghai Yellow chicken. A total of 1,300,074,528 clean reads and 27,023 lncRNAs were obtained from twenty samples. 3095 genes (1,336 lncRNAs and 1,759 mRNAs) were differentially expressed among different stages, of which the number of DEGs decreased with the differentiation, demonstrating that the early stage might be most important for chicken preadipocytes differentiation. Furthermore, 3,095 DEGs were clustered into 8 clusters with their expression patterns by K-means clustering. We identified six stage-specific modules related to A0, A2 and A6 stages using weighted co-expression network analysis. Many well-known/novel pathways associated with preadipocytes differentiation were found. We also identified highly connected genes in each module and visualized them by cytoscape. Many well-known genes related to preadipocytes differentiation were found such as *IGFBP2* and *JUN*. Yet, the majority of high connected genes were unknown in chicken preadipocytes. This study provides a valuable resource for chicken lncRNA study and contributes to batter understanding the biology of preadipocytes differentiation in chicken.

## 1. Introduction

The abdominal fat is an important carcass trait of broiler. The production performance of broiler has been significantly improved after decades of breeding and selection. However, the overemphasis on selection of broilers for rapid growth rate leads to excessive fat accumulation, especially for Chinese local chicken breeds. The excessive fat is often disposed as waste. Furthermore, excess fat deposition results in reduction of feed conversion ratio, carcass yield, laying rate, fertility rate and hatching rate. Lower abdominal fat therefore has become one of the breeding goals of broiler.

Adipose tissue is a complex, essential, and highly active metabolic and endocrine organ(Kershaw and Flier, 2004). The growth of adipose tissue is primarily due to the increase in adipocyte cell number (hyperplasia) and the enlargement of adipocytes (hypertrophy). As is well known that adipocytes are derived from pluripotent mesenchymal stem cells (MSCs) in adipogenesis process(Ding et al., 2015). The MSC have ability to develop into adipoblasts, which then will develop into preadipocytes that could store lipid. The preadipocytes can finally differentiate into adipocytes under particular conditions(Leclercq, 1984). The cell number in mature adipose tissue is thought to reflect the proliferation of preadipocytes and their subsequent differentiation into mature adipocytes(Matsubara et al., 2013).

The adipogenesis process is controlled by a complex process that regulated by various transcriptional events. In mammals, the differentiation of preadipocytes is well studied, especially in human and mouse. Previous studies have identified that peroxisome proliferator-activated receptor γ (*PPARγ*) and CCAAT/enhancer binding proteins (*C/EBPs*) are key genes regulating adipocytes differentiation(MacDougald and Lane, 1995; Morrison and Farmer, 2000). Matsubara et al. reported that PPARγ is also a key regulator of preadipocyte differentiation(Matsubara et al., 2005). Rencent studies in mammals have demonstrated functions of some new transcription factors in adipogenesis, such as transcriptional factor of zinc finger protein 423 (*Zfp423*)(Gupta et al., 2010), Krüppel-like transcription factors (KLFs) (*FGF10*)(Banerjee et al., 2003; Kaczynski et al., 2003; Mori et al., 2005) and Fibroblast growth factor 10(Sakaue et al., 2002; Yamasaki et al., 1999).

In chicken, several genes including *KLF2*(Zhang et al., 2014a), *KLF3*(Zhang et al., 2014b) and *FATP1*(Qi et al., 2013) are identified as regulator in chicken adipogenesis and preadipocytes differentiation. However, little is known about the details of how adipogenesis is regulated. Recently, several studies try to investigate the regulation mechanism of chicken adipogenesis by genome-wide analysis of mRNA(Ji et al., 2012; Regassa and Kim, 2015) and microRNA(Wang et al., 2015). In the present study, lncRNA and mRNA profiling of preadipocytes during differentiation were analyzed using RNA sequencing. Our study focused on characterizing the features of lncRNA and identifying differentially expressed lncRNAs and mRNAs between different differentiation stages of preadipocytes. The functions of differentially expressed genes were annotated and the pathways involved were enriched. Our study provides a valuable resource for chicken lncRNA study and contributes to batter understanding the biology of preadipocytes differentiation.

## 2 Materials and mathods

### 2.1 Primary culture of chicken preadipocytes from abdominal adipose tissue

Chicken preadipocytes from abdominal adipose tissue were cultured according to the method described by Shang(Shang et al., 2014), with some modifications. Abdominal adipose tissue was collected from 14-day-old Jinghai yellow chicken under sterile condition. Adipose tissue was washed by PBS supplemented with penicillin (100 units/ml) and streptomycin (100 μg/ml). The washed tissue was cut to 1 mm 3 by surgical scissors and then digested using 2 mg/ml collagenase type I (Sangon Biotech) with shaking for 65 min at 37 °C. The digested cell suspension was filtrated using 200 and 500 mesh screens and centrifuged at 300g for 10min (22 °C) to separate the stromal-vascular fractions from undigested tissue debris and mature adipocytes. Stromal-vascular cells were plated to 50mm culture palte at a density of 1×105 cells/ml and cultured with DMEM/F12 (Dulbecco’s modified Eagle’s medium/Ham’s nutrient mixture F-12) basic medium (10% (v/v) FBS, 100 units/ml penicillin and 100 μg/ml streptomycin) in a humidified atmosphere with 5% (v/v) CO2 at 37 °C until reaching 90% confluence.

### 2.2 Induction of abdominal preadipocytes

Following 90% cell confluence, the basic medium was removed and replaced with differentiation medium (0.25 μM dexamethasone (Takara), 10 μg/ml insulin (Takara) and 0.5 mM IBMX (Takara) for 48 hours. The differentiation medium was replaced with maintenance medium (10 μg/ml insulin (Takara)) and incubated for 48 hours. The detailed procedure for induction of abdominal preadipocytes was described in Figure 1. Cells were collected after induced for 0h, 48h, 96h and 144h. Each point included 3 biological replicates (n=3).

**Figure 1.**
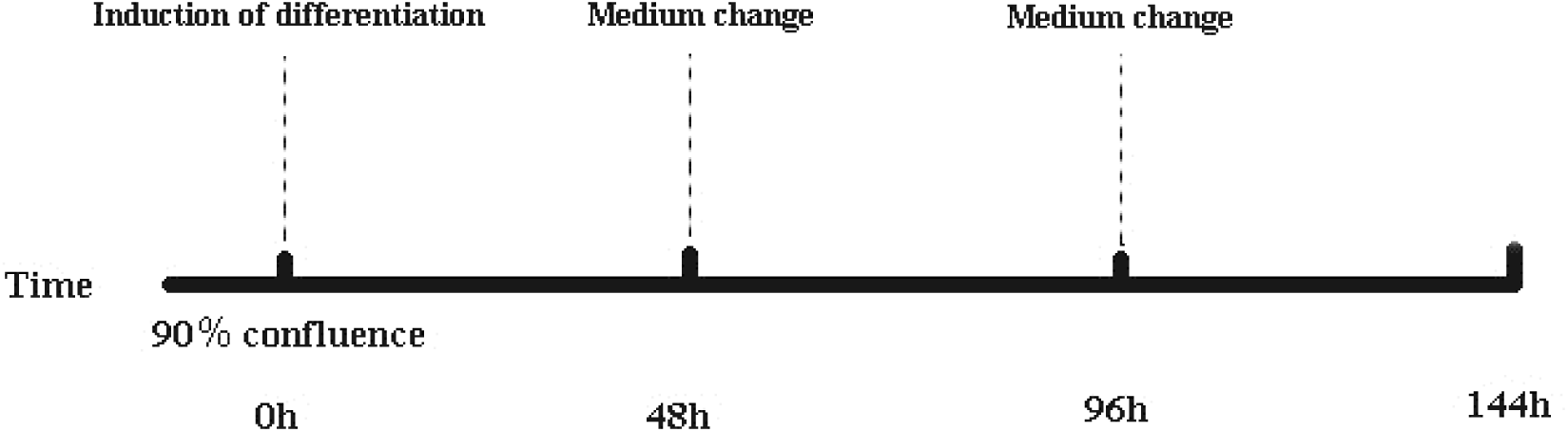
Induction differentiation procedure of abdominal preadipocytes. The basic medium consisted of DMEM/F12 and 10% FBS. Induction differentiation medium consisted of basic medium, insulin, dexamethasone and IBMX. Maintenance medium consisted of basic medium and insulin. Induction differentiation medium was replaced with maintenance medium at 48h, while maintenance medium was replaced with basic medium at 96h.

### 2.3 RNA extraction, library construction and sequencing

A total of 12 cell samples were successfully collected. Total RNA was extracted using Trizol reagent (Invitrogen). The integrity, concentration and purity of total RNA were checked via Nanodrop, Qubit 2.0 and Aglient 2100. RNA samples with a RIN value greater than 8.0 and an OD 260/280 ratio greater than 1.9 were selected for deep sequencing. The rRNA was removed and mRNA was enriched using magnetic bead with Oligo (dT) and then randomly fragmented by Fragmentation Buffer. The mRNA was used as template to synthesize the first-strand cDNA using 1st Strand Enzyme Mix (Enzyme). The second-strand cDNA was synthesized using 2nd Strand Marking Buffer and 2nd Strand/End Repair Enzyme Mix (Enzyme). The products were purified by VAHTS™ DNA Clean Beads and then the end of double strand was repaired and A-tailed. An adapter was jointed to A-tailed products using ligation mix. Suitable sized fragments were selected using VAHTS™ DNA Clean Beads to construct the cDNA library by PCR. The RNA sequencing was performed using Illumina HiSeq2500.

### 2.4 Quality control

The raw data was performed to quality control using FastQC (http://www.bioinformatics.babraham.ac.uk/projects/fastqc/). The base composition and quality distribution of reads and the GC and AT base content were analyzed, which could reflect the quality of raw data in whole. Clean Data was obtained by removing reads containing adapter, reads containing over 10% of ploy-N, and low-quality reads (>50% of bases whose Q scores were ≤10%) from the raw data.

### 2.5 Sequencing data analysis and transcriptome assembling

Clean Data was mapped to the *Gallus gallus* reference genome (gal4) by Tophat(Trapnell et al., 2012) program using the following parameters: --segment-length 25 –segment-mismatches 2. The remaining parameters were set as default. The uniformity, insert length and saturation of sequencing data were analyzed based on the alignment results. The transcripts were assembled using cufflinks(Trapnell et al., 2012) program based on RABT Assembling strategies.

### 2.6 lncRNA prediction

Based on the assembling results, transcripts with RPKM=0 were wiped off. The filter criteria of lncRNA were: 1) removed transcripts shorter than 200nt; 2) removed transcripts with a ORF that was longer than 300nt; 3) removed transcripts containing specific domain; 4) transcripts that were similar to known protein; 5) transcripts that were predicted to coding by CPC.

### 2.7 lncRNA targets prediction and annotation

lncRNA functioned by acting on protein coding gene via *cis*-acting element and trans-acting factor. In the present study, lncRNA targets were predicted based on *cis* function prediction. The closest coding genes to lncRNAs in 10K of upstream and downstream were screened and their associations with lncRNA were analyzed using Bedtools program(Quinlan and Hall, 2010). Then the target genes were conducted to functional enrichment analysis using the DAVID database(Huang da et al., 2009).

### 2.8 Quantitation of gene expression

Cuffdiff(Trapnell et al., 2013) was used to calculate reads per kb for a million reads (RKPM) of both mRNA and lncRNA in each sample. For biological replicates, transcripts or genes with a *p*<0.05 and foldchange≥2 were defined as differential expressed genes or lncRNAs between two groups(Ren et al., 2016).

### 2.9 Co-expression network analysis

The co-expression network was contructed by Weighted Gene Co-expression Network Analysis (WGCNA) package(Langfelder and Horvath, 2008) in R environment with the 3,095 differentially expressed genes. Module was detected by dynamic tree cutting method. Then the stage-specific modules were identified based on the correlation between eigengene and traits. Module with a *P*-Value <0.05 was defined as significant. We identified the central and highly connected genes by visualizing the top 200 connections of the top 150 genes for each stage-specific module.

### 2.10 Gene ontology and Kyoto Encyclopedia of Genes and Genomes analysis

Functional annotation enrichment analysis for Gene Ontology (GO) and Kyoto Encyclopedia of Genes and Genomes (KEGG) were conducted by DAVID database(Huang da et al., 2009). Go terms and pathways with *P* value less than 0.05 were considered as significantly enriched.

### 2.11 Validation of gene and lncRNA expression by qRT-PCR

Primers were designed using the Primer-BLAST in NCBI (http://www.ncbi.nlm.nih.gov/tools/primer-blast/) (Table 1). The first cDNA strand was synthesized using PrimeScript™ RT Master Mix (Perfect Real Time) kit (Takara, JPN) according to the user manual. β-actin was used as housekeeping gene. Expression of lncRNA and mRNA were quantified using SYBR® Premix Ex Taq™ II kit (Takara, JPN). The 20 μL PCR reaction included 10 μL SYBR Premix Ex Taq II (TaKaRa, Dalian,China), 0.4 μL (10 p moL / μL) specific forward primer, 0.4μL (10 p moL /μL) reverse primer, 0.4 μL ROX reference dye, 2 μL (10 ng/μL) diluted cDNA and 6.8 μL RNase free water. Cycling parameters were 95°C for 30 s, followed by 40 cycles of 95°C for 5 sec and 60°C for 34 sec. Melting curve analyses were performed following amplifications. The ABI 7500 software was used to detect the fluorescent signals. Quantification of selected gene expression was performed using the comparative threshold cycle (2^−ΔΔCT^) method.

**Table 1.**
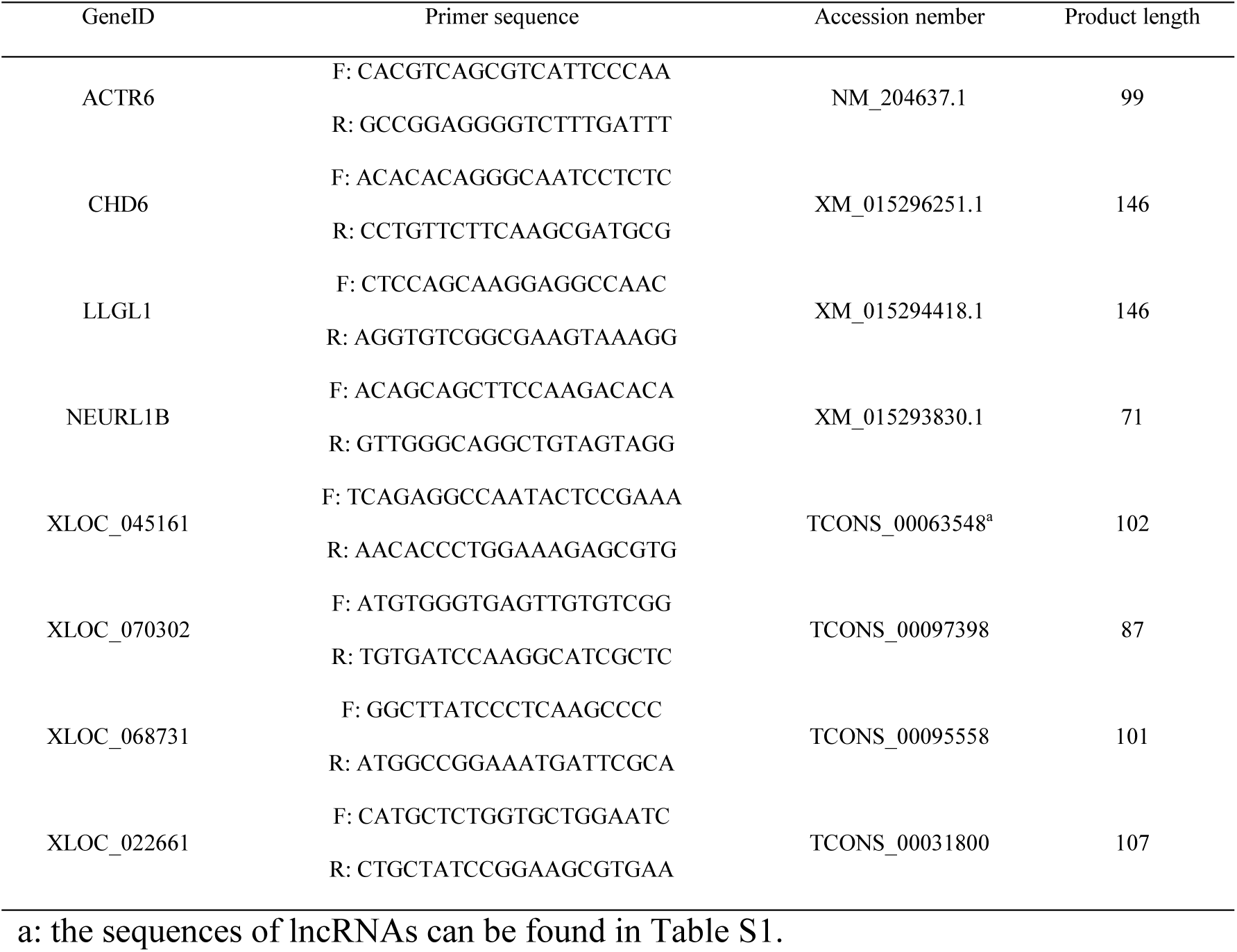
Primers of genes for qRT-PCR

**Table 1.**
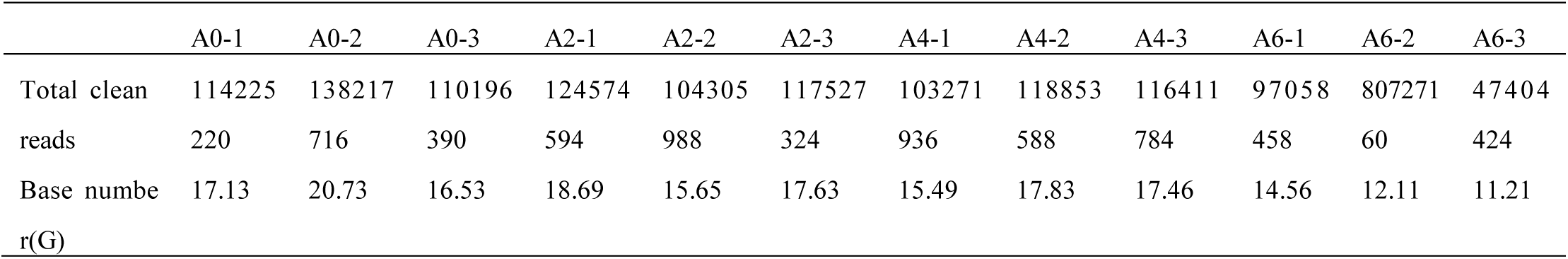
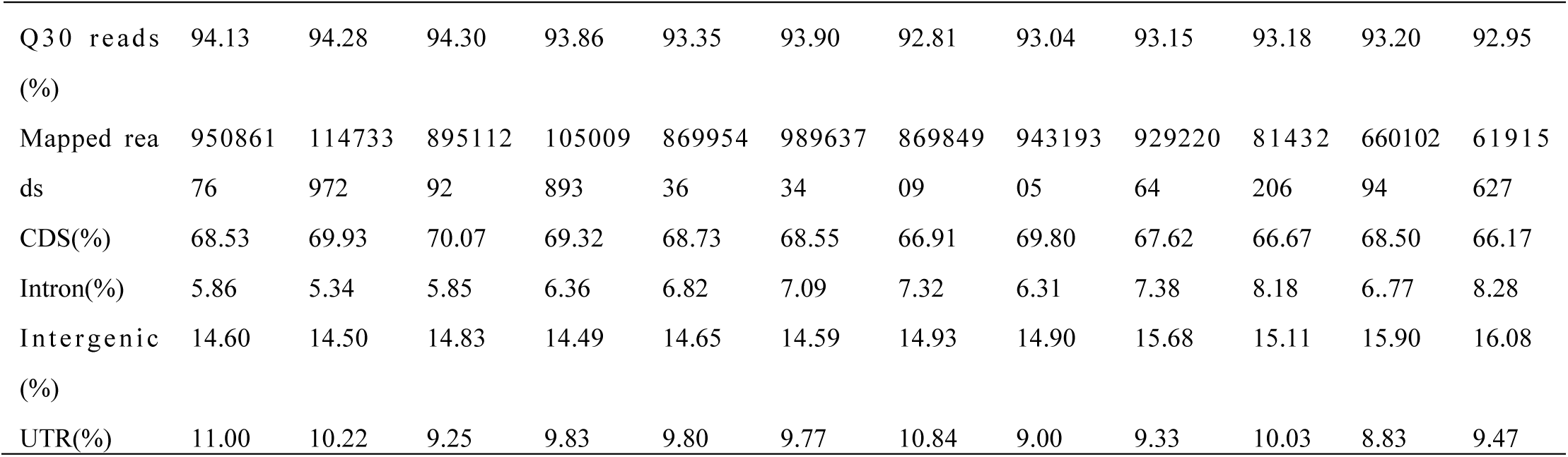
Statistics of clean reads in preadipocytes of chicken

The Sequencing data of our study were submitted to the sequence Read Archive (Accession Number SRR3985377) in NCBI. Supplemental materials include ten files. Table S1 contains the names and sequences of all the candidate lncRNAs. Table S2 contains all the differentially expressed lncRNAs and mRNAs. The common differentially expressed lncRNAs and mRNAs among three comparisons (A0 vs A2, A2 vs A4, A4 vs A6) are included in Table S3. Table S4, Table S5 and Table S6 are the GO and pathway analysis of target genes of lncRNAs, DEGs of different stages and co-expressed mRNAs with lncRNAs, respectively. Table S7, Table S8 and Table S9 are the annotation of genes in A0, A2 and A6 stage-specific modules, respectively. The validation of RNA-seq using qRT-PCR is included in Table S10. Figure S1 and Figure S2 are GO analysis of DEGs of different stages and visualization of the co-expression network of all DEGs.

## 3. Results

### 3.1 Sequencing results and quality control

A total of 1,394,219,096 raw reads were produced from 12 cDNA libraries. After quality control, 1,300,074,582 clean reads (195.02 Gb) were obtained. The percentage of clean reads among raw reads in each library ranged from 91.41% to 94.73%. The percentage of reads with a Phred quality value greater than 30 among clean reads ranged from 92.95% to 94.30%. The average GC content of clean reads in 12 samples was 52.51%. Subsequently, the clean reads were aligned with the chicken reference genome (http://hgdownload.soe.ucsc.edu/goldenPath/galGal4/bigZips/galGal4.fa.gz). The mapped rate of 12 samples ranged from 79.40% to 84.30%. Among these mapped reads, 66.17%-70.07% of reads were mapped to CDS regions, 5.34%-8.18% to intron regions, 14.49%-16.08% to intergenic regions, and 8.83%-11.00% to UTR regions (Table 2). High Pearson correlation coefficients were found among biological replicates of the same differentiation stage, which indicated the reproducibility of sample preparation (Figure 5A).

### 3.2 Identification of lncRNAs in abdominal preadipocytes

We used five tools, namely NONCODE, TransDecoder, Pfam, BlastX and CPC, to remove potential coding and short (length<200nt) transcripts. Finally, 27,023lncRNAs were obtained (Table S1). The length and exon number of lncRNAs were analyzed. We found that lncRNAs were shorter in length and fewer in exon number than protein coding genes in abdominal preadipocytes (Figure 2).

**Figure 2.**
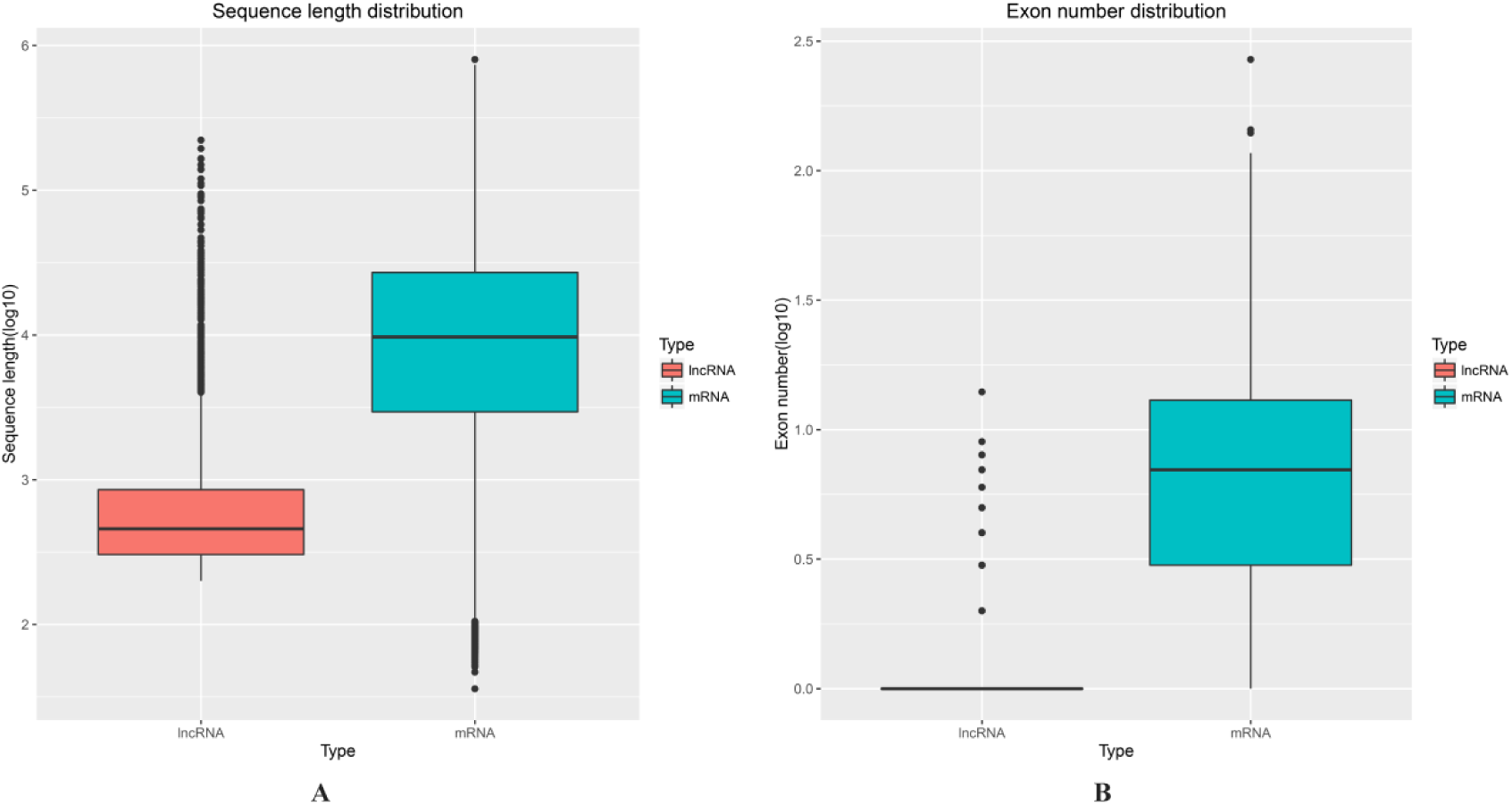
Length and exon number of lncRNAs and mRNAs. (A) Length distributionog mRNAs and lncRNAs. (B) Exon number distribution of mRNAs and lncRNAs.

### 3.3 Differentially expressed lncRNAs and mRNA during abdominal fat preadipocytes differentiation

Given a *P* value < 0.05 and Fold change≥2, 1,336 differentially expressed lncRNAs and 1,759 differentially expressed mRNAs were obtained by pairwise comparison (A0 vs A2; A0 vs A4; A0 vs A6; A2 vs A4; A2 vs A6 and A4 vs A6) of samples collected from preadipocytes at day 0, 2, 4 and 6 of differentiation (Figure 3A, 3B) (Table S2). No common gene was identified among 6 comparisons. 936 differentially expressed lncRNAs and 1,280 differentially expressed mRNA were obtained by pairwise comparison (A0 vs A2; A2 vs A4; A4 vs A6) of the same samples. As shown in figure3C, D, 30 differentially genes (DEGs) were common among three comparisons (7 lncRNAs and 23 mRNAs) (Table S3). We counted the DEGs number of three comparisons (A0 vs A2, A2 vs A4 and A4 vs A6) (Figure 3E, F). The results showed that the number of differentially expressed mRNAs and lncRNAs decreased with the differentiation of preadipocytes.

**Figure 3.**
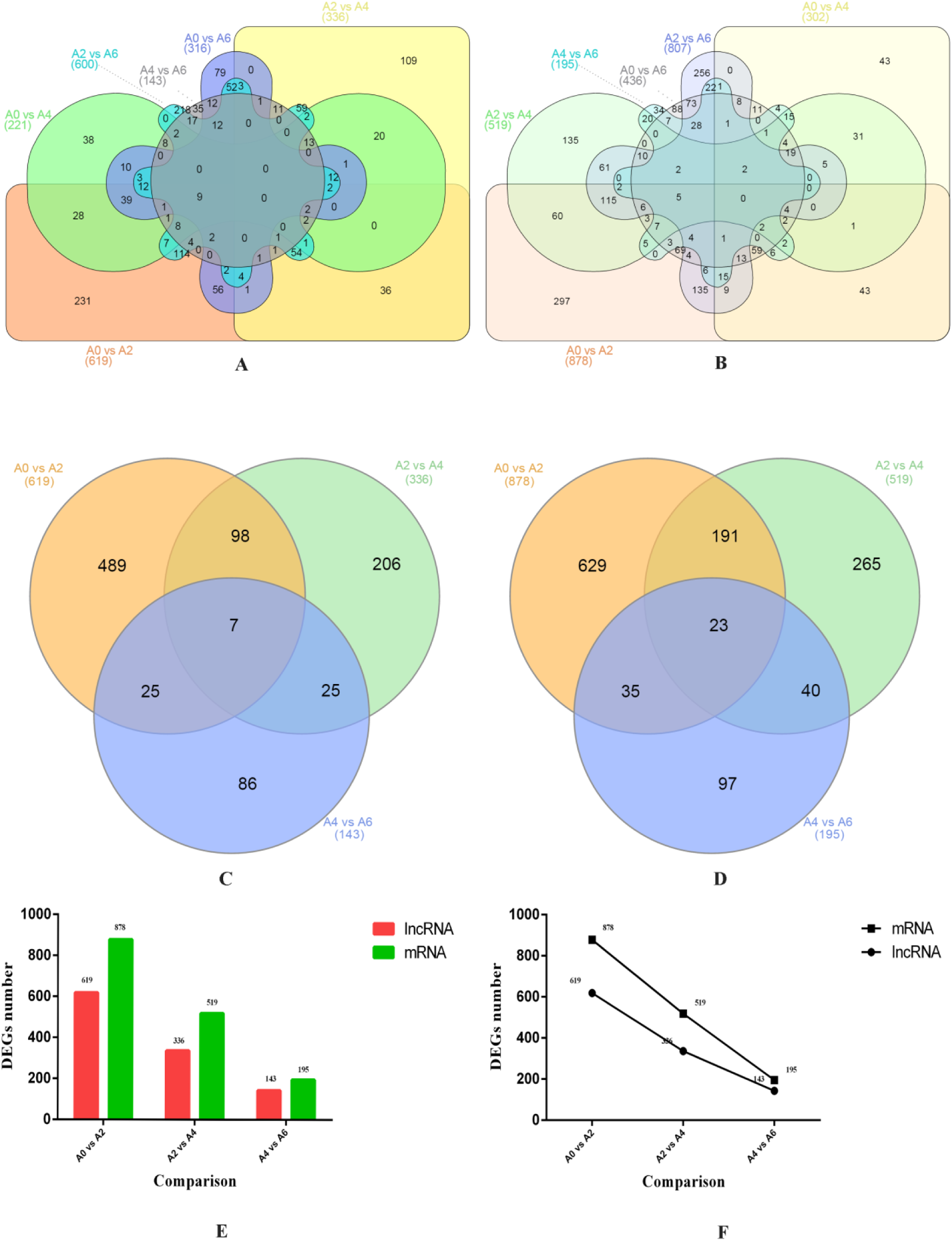
Venn diagram of DEGs at different stages. (A) Venn diagram of common LncRNAs in six comparison (A0 vs A2, A0 vs A4, A0 vs A6, A2 vs A4, A2 vs A6 and A4 vs A6). (B) Venn diagram of common mRNAs in six comparison (A0 vs A2, A0 vs A4, A0 vs A6, A2 vs A4, A2 vs A6 and A4 vs A6). (C) Venn diagram of common LncRNAs in three comparisons (A0 vs A2, A2 vs A4 and A4 vs A6). (D) Venn diagram of common mRNAs in three comparisons (A0 vs A2, A2 vs A4 and A4 vs A6). (E) Histogram of DEGs number in three comparisons. (F) Line chart of DEGs number in three comparisons.

We performed K-means clustering of all DEGs using the euclidean distance method associated with complete linkage (Figure 4). Eight clusters were plotted with expression patterns (Table S4). The K1 cluster included 392 genes which showed up-regulated at day two of differentiation and then down-regulated at day 4 and 6 of differentiation. The 49 genes in K2 were up-regulated across the whole induction process. The expressions of 159 genes in K3 were slightly down-regulated at A4 and A6 stage compared to A0 and A2 stage. K5 cluster included most of the DEGs which expressed smoothly across four stages proceeded. The genes in K5 and K5 clusters had opposite expression pattern. The genes in K5 were significantly down-regulated at A2, A4 and A6 stages compared to the first stage, whereas, the genes in K6 were significantly up-regulated at A2, A4 and A6 stages compared to the first stage. The expression level of genes in K7 remained stable at first two stages and then was up-regulated at last two stages.

**Figure 4.**
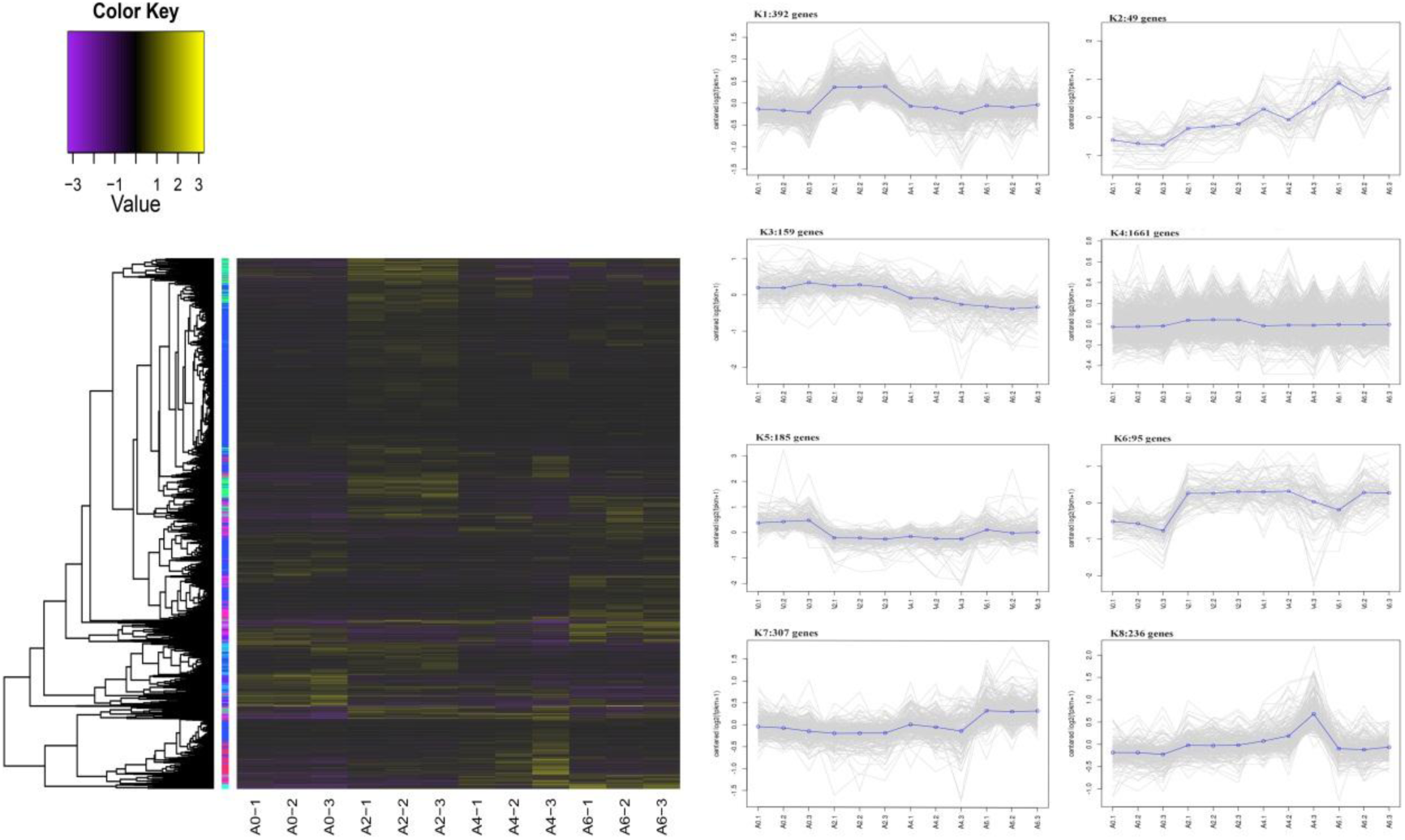
Clustering of all DEGs (lncRNAs and mRNAs). (A) The heat map shows the K-means clustering of transformaed expression values for lncRNAs and mRNAs. Yellow represents higher expression and purple represents lower expression. (B) expression patterns of genes in eight clusters corresponding to the heat map.

### 3.4 Functional prediction of lncRNAs and mRNAs in preadipocytes samples

We selected coding genes 10K upstream and downstream of lncRNAs as the *cis* target genes(Ren et al., 2016; Wang et al., 2016). Finally, 4915 target genes were identified. To predict the function of lncRNAs in preadipocytes of chicken, we performed GO and KEGG analysis with the *cis* target genes (Table S4). A total of 1746, 1544 and 2174 genes were assigned to biological process, cellular component and molecular function GO categories, respectively. In the biological process category, 27 terms such as cellular process, system development and anatomical structure development were significantly enriched. In cellular component category, the top three terms were intracellular, intracellular membrane-bounded organelle and membrane-bounded organelle, while in molecular function category, protein binding and binding protein serine/threonine kinase activity were the most abundant terms. The KEGG enrichment showed that 971 out of 4915 genes were significantly enriched in 9 pathways including Wnt signaling pathway, MAPK signaling pathway and Vascular smooth muscle contraction pathway.

To gain insight into the similarities and differences in differentiation of three stages, the differential expressed mRNAs of three comparisons (A0 vs A2, A2 vs A4, A4 vs A6) were conducted to GO and KEGG pathway analysis (Table S5). In the biological process category, most of the genes were involved in processes associated with cellular regulation and metabolism, such as cellular process, cellular macromolecule metabolic process and cellular metabolic process. The top three GO terms were cellular process, metabolic process and primary metabolic process at A0-A2 and A4-A6 stages, while they were biological regulation, regulation of biological process and regulation of cellular process at A2-A4 stage (Figure S1). KEGG analysis showed that, in the top fifteen pathways, five pathways were common among three stages. They were Cell cycle, Cytokine-cytokine receptor interaction, Jak-STAT signaling pathway, Oocyte meiosis and Ubiquitin mediated proteolysis pathways. Furthermore, 5, 4 and 5 stage-specific pathways were identified for three differentiation stages. Stage-specific pathways with the highest number of DEGs for A0-A2, A2-A4 and A4-A6 stages were Oxidative phosphorylation, Regulation of actin cytoskeleton and Glycosphingolipid biosynthesis, respectively.

### 3.5 Co-expression network and module construction

In our study, WGCNA R software package was used to perform the co-expression network analysis. lncRNAs exerted its biological functions by regulating target mRNAs. The co-expression network could help us to predict target mRNAs of lncRNAs, which shared common expression patterns. To gain insight in the functions of differential expressed lncRNAs, we constructed co-expression network with 1336 and 1759 differential expressed lncRNAs and mRNAs (Figure S2). The node and edge files respond to gene expression profiles and pairwise correlations between gene expressions, respectively. 106 mRNAs were identified to have common expression patterns with 176 lncRNAs, which might be target genes of lncRNAs (Table S6). The 106 mRNAs were conducted to GO and KEGG analysis to predict the functions of lncRNAs. GO analysis showed that signal transduction, regulation of biological process and system development were the top three terms in biological process category. In molecular function category, insulin-like growth factor binding, nucleoside-triphosphatase activity and pyrophosphatase activity terms were significantly enriched while in cellular component, trans-Golgi network transport vesicle was significantly enriched. No significant pathway was enriched. However, several well-known pathways involved in differentiation of preadipocytes were identified, including ErbB, MAPK, Insulin and Jak-STAT signaling pathway.

Weighted gene co-expression network analysis can be used for finding clusters (modules) of highly correlated genes, for summarizing such clusters using the module eigengene or an intramodular hub gene, for relating modules to one another and to external sample traits(Langfelder and Horvath, 2008). In our study, 3,095 DEGs were used to identify groups of co-expressed genes, termed “modules”. Each module is assigned a unique color label underneath the cluster tree(de Jong et al., 2012). Ten modules were identified in size ranging from 99 genes in the purple module to 524 in the blue module (Figure 5B). The co-expression modules could not exist independent, instead, they formed a meta-network. To explore and identify the correlations among modules, the modules were conducted to clustering analysis based on their eigengenes. The results showed that ten modules were classified into 4 groups: the first was turquoise module; the second included magenta, brown and pink modules; the third included purple and red modules; the fourth included blue, black, green and yellow modules (Figure 5C). Modules classified into same group might have same or similar functions and regulation mechanisms.

**Figure 5.**
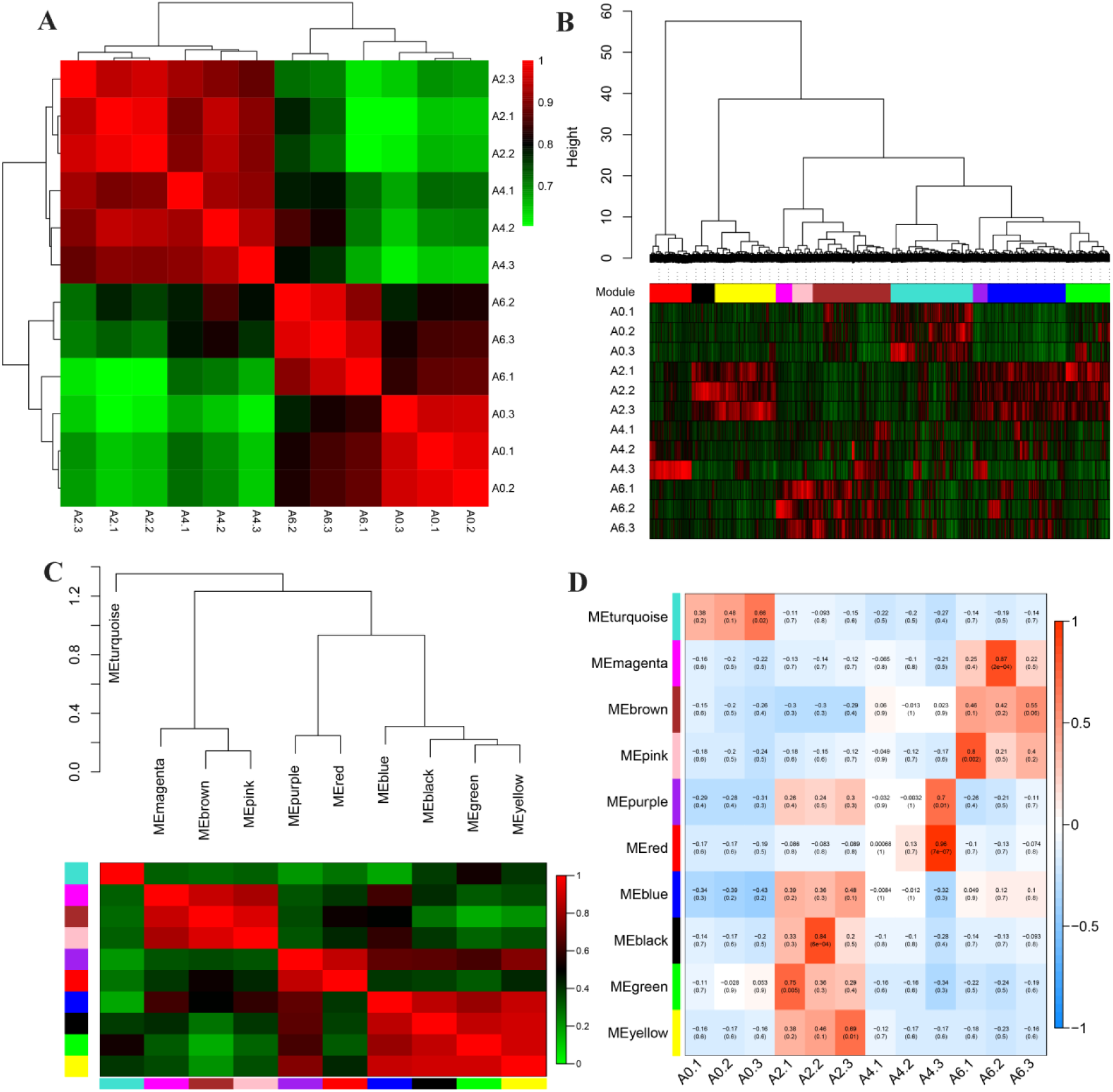
Visualization of co-expression network construction. (A) Heatmap of duplicate samples of chicken abdominal preadipocytes differentiation. The color, ranging from green through dark to red, represents pearson correlation coefficients ranging from 0.6 to 1, indicating low to high correlations. All samples in the same differentiation stage are highly correlated in pearson coefficients indicating the reproducibility of the samples. (B) Hierarchical cluster tree (average linkage, dissTOM) of the 3095 genes. The color bands provide a simple visual comparison of module assignments (branch cuttings) based on the dynamic hybrid branch cutting method. (C) Clustering of modules based on eigengenes. The color, ranging from green through dark to red, represents pearson correlation coefficients ranging from 0 to 1, indicating low to high correlations. (D) Heatmap of correlations between module and differentiation stage. The color, ranging from blue through white to red, indicates low to high correlations.

### 3.6 Identification and visualization of stage-specific module

To explore stage-specific modules during chicken abdominal preadipocytes differentiation development, we calculated the *gene significance* (GS) and *module membership* (MM) of all genes within a module. GS was defined as (the absolute value of) the correlation between the gene and the differentiation stage. MM was defined as the correlation of the module eigengene and the gene expression profile. We identified six stage-specific modules (FDR *P*<0.05) (Figure 6), in which the black, blue, green and yellow modules were positively correlated with A2 stage (Figure 6A, B, D, F) while the turquoise and brown modules were positively correlated with A0 and A6 stages (Fig 6C, E), respectively. It meant that the genes in modules above were predominantly up-expressed at day 2, day 0 and 6 of differentiation. Furthermore, the blue module also was negatively correlated with A0 stage, meaning that the genes in blue module were predominantly down-expressed at day 0 of differentiation. We also found that the magenta, pink and purple modules, though not significant (FDR *P*>0.05), might play important role in preadipocytes differentiation when it comes to their expression pattern (Figure 5B, D). Genes in magenta and pink modules expressed in low level before day 4 of differentiation and then achieved a relatively high expression level at day 6 of expression. Similarly, the genes in purple module expressed low at day 0, increased their expression level at day 2 and then decreased their expression level at day 4 and 6 of differentiation.

**Figure 6.**
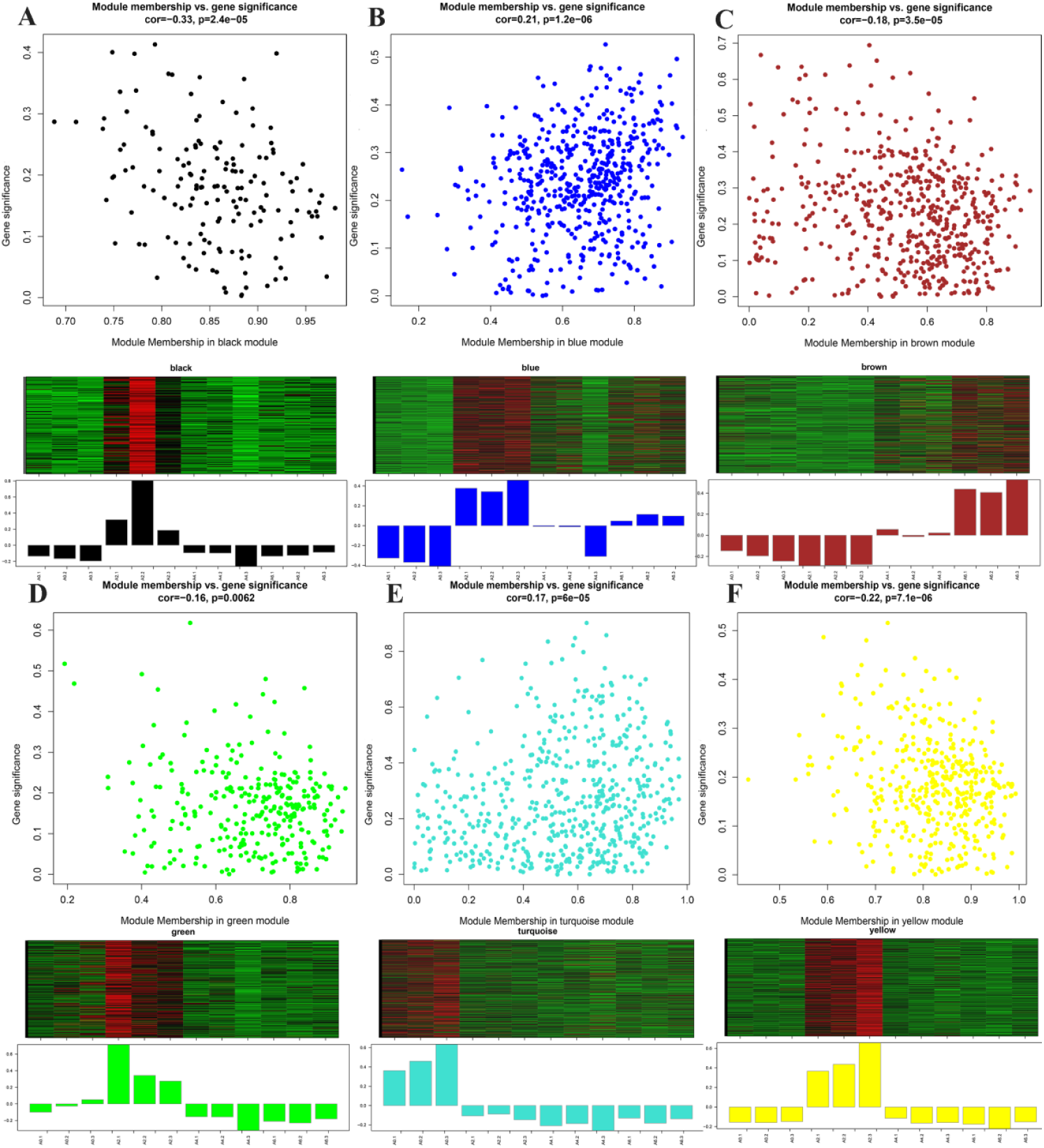
visualization of GS vs MM and gene expression level of significant modules. Scatterplot represents the GS and MM of each module, which exhibits a significant correlation between GS and MM (*P*<0.05), implying that module tend to be associated with differentiation stage. Clustering heatmap and barplot represent gene expression level of each module. In the heatmap, the color range from green through dark to red indicates low to high expression level.

### 3.7 identification of central and high connected genes

In order to identify genes that are central and highly connected in the stage-specific modules, we analyzed top 200 connections of the top 150 highly connected genes for each stage-specific module and visualized them using Cytoscape (Figure 7). Five common genes (Three lncRNAs and two mRNAs) were identified among four highly correlated modules of black, blue, green and yellow. It suggested that XLOC_068731, XLOC_022661, chromodomain helicase DNA binding protein 6 (*CHD6*), lethal giant larvae homolog 1 (*LLGL1*) and neuralized E3 ubiquitin protein ligase 1B (*NEURL1B*) might play important roles in differentiation of A2 stage. In the brown module, Kelch like family member 38 (*KLHL38*) and XLOC_045161 were the highest connected mRNA and lncRNA, indicating their potential key roles in regulation of preadipocytes differentiation at A6 stage. ARP6 actin-related protein 6 homolog (*ACTR6*) and XLOC_070302, with the most connection, were found to be important factors in A0 stage of differentiation.

**Figure 7.**
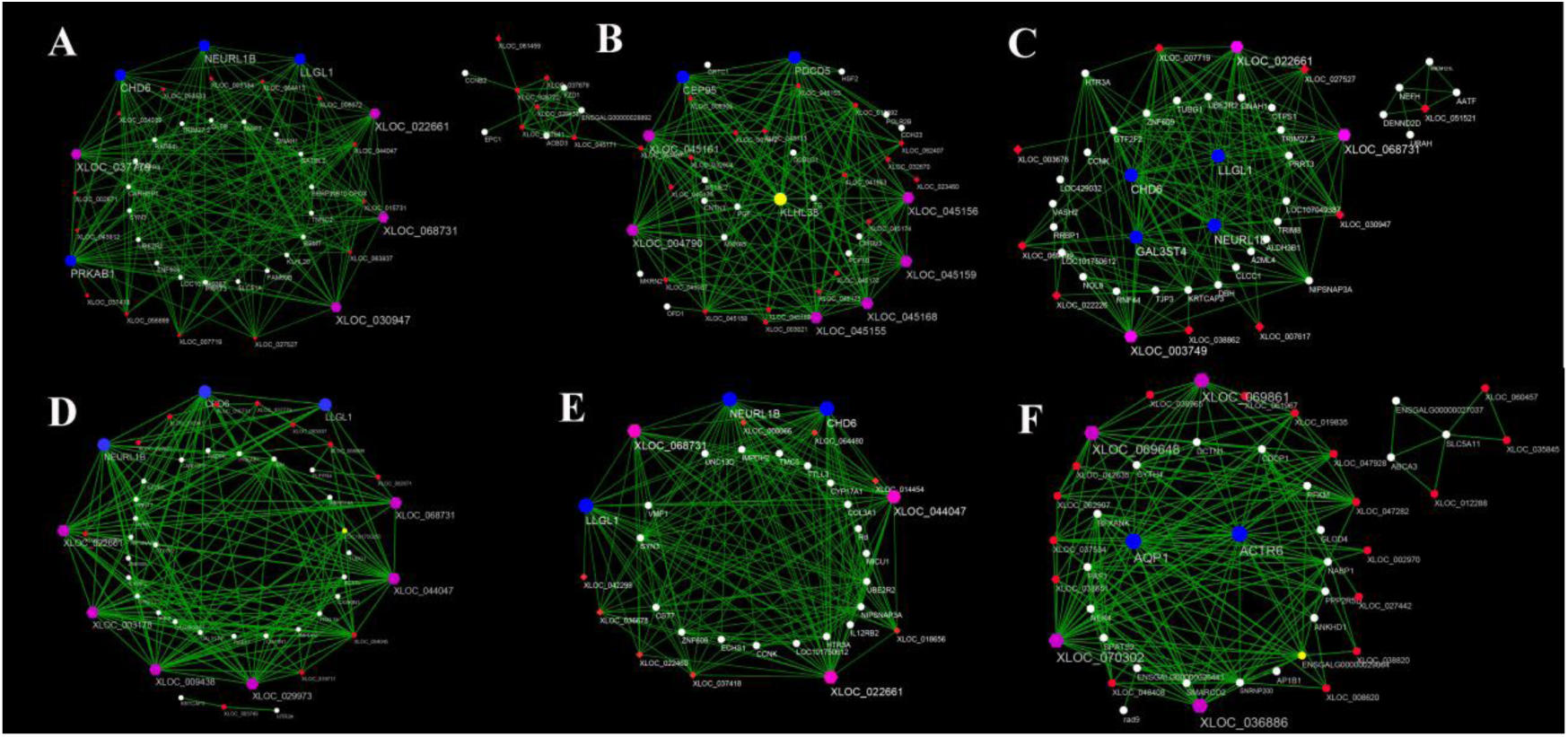
visualization of connections of genes in modules. A, B, C, D, E and F represent the visualization of connections of genes in black, brown, green, yellow, blue and turquoise modules. Red-colored nodes represent mRNAs. White-colored nodes represent lncRNAs. Pink-colored and blue-colored nodes indicate common and highly connected lncRNAs and mRNAs, suggesting their central role in the network.

### 3.8 GO and pathway analysis of genes in stage-specific modules

GO analysis reveals the functions of genes in stage-specific modules and pathway analysis reveals essential signaling and metabolic networks in preadipocytes differentiation. The enriched GO terms of biological process and pathways were showed in Table S7, S8 and S9. For GO analysis, RNA metabolic process, regulation of localization and cellular metabolic process were significantly enriched at A0 stage, generation of precursor metabolites and energy, electron transport chain and energy derivation by oxidation of organic compounds were highly enriched at A2 stage, while blastocyst development and cellular process were greatly enriched at A6 stage (*P*<0.05). In the pathway analysis, no significant pathway was identified at A0 and A6 stages. Oocyte meiosis, Lysosome and Galactose metabolism were the top three pathways at A0 stage, while Biosynthesis of unsaturated fatty acids, Apoptosis and Propanoate metabolism were top three pathways at A6 stage. When it came to A2 stage, Oxidative phosphorylation was significantly enriched (P<0.05). Interestingly, three common pathways for Oxidative phosphorylation, Lysosome and Propanoate metabolism were found in different differentiation stages. Additionally, many well-known pathways related to preadipocytes differentiation including ABC transporters, Jak-STAT, Wnt, Insulin, Fatty acid metabolism and Fatty acid biosynthesis signaling pathways were found.

### 3.9 Validation of DEGs by qRT-PCR

Quantitative real time PCR (qRT-PCR) was carried out to validate the central and highly connected genes XLOC_045161, XLOC_070302, XLOC_068731, XLOC_022661, *ACTR6*, *CHD6*, *LLGL1* and *NEURL1B*. We used the same 12 cell samples as were used in the RNA-seq for qRT-PCR validation. The qRT-PCR results for all the genes were tested statistically by the T-test method. The results showed that the expression patterns of these 8 genes were in excellent agreement with the RNA-seq results(Figure 8, Table S10).

**Figure 8.**
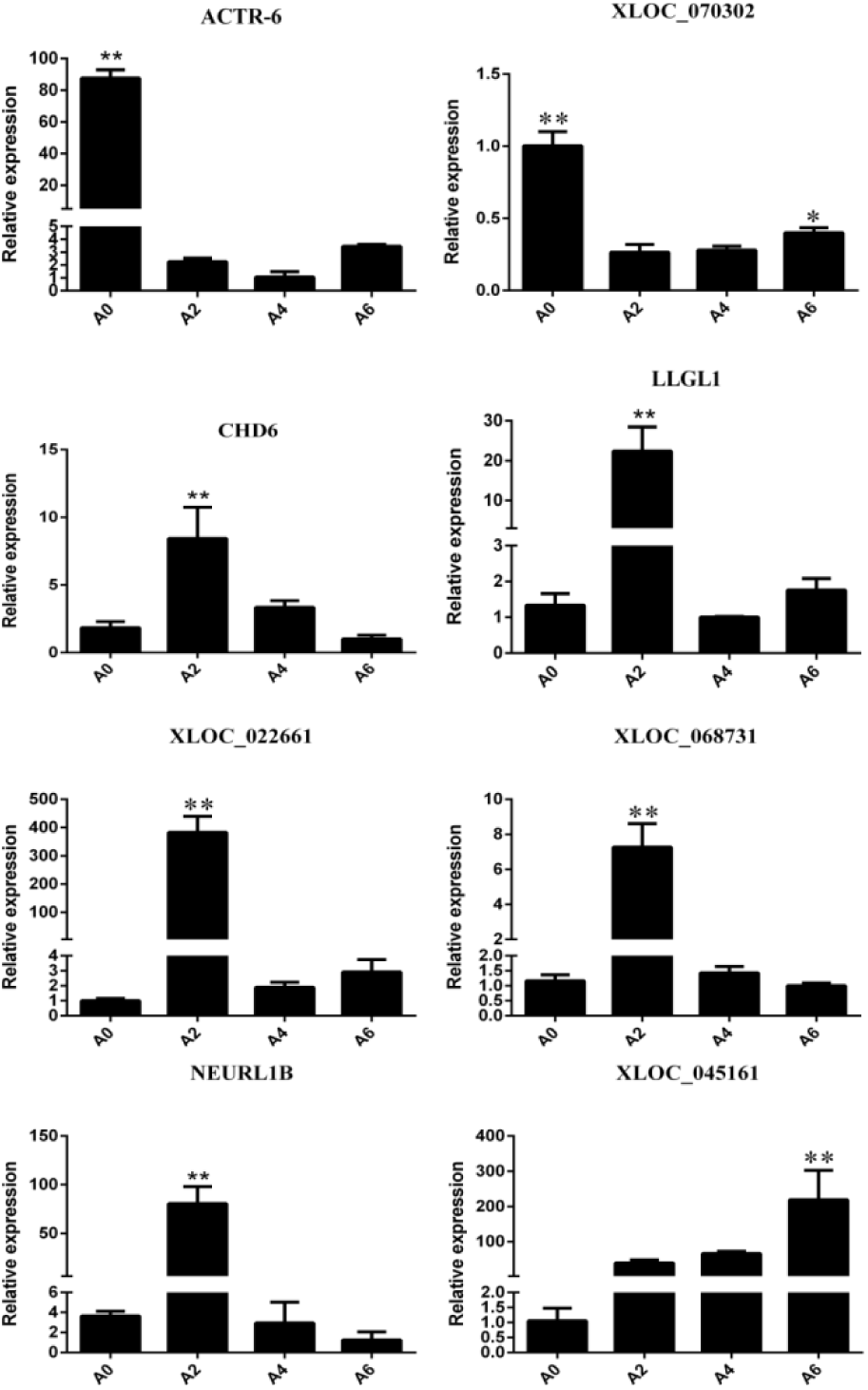
Validation of highly connected genes using qRT-PCR

## 4. Discussion

In the last 60 years, the selection of important economic traits has been the focus and significant genetic improvements have been gained(Dong et al., 2015). Genetic selection for enhanced growth rate in meat-type chickens (Gallus domesticus) is usually accompanied by excessive adiposity, which has negative impacts on both feed efficiency and carcass quality(Resnyk et al., 2015). The genetic improvement of abdominal fat by standard selection has been lower for two reasons: (1) intensity of selection has been lower because of the difficulty and cost to measure these phenotypes, and (2) genetic evaluations have been less accurate because evaluation of candidates is based on information from relatives only(Demeure et al., 2013). Exploring the molecular regulation mechanisms of fat deposition in broilers is helpful for optimal breeding. Many studies have been carried out on gene expression of chicken abdominal fat using genome-wide RNA sequence(Resnyk et al., 2013; Resnyk et al., 2015; Zhuo et al., 2015).

Long non-coding RNA (lncRNA) is a kind of non-coding RNA longer than 200 nt, which has attracted much attentions in the past several years. Studies show that lncRNAs regulate metabolic tissue development and function, including adipogenesis, hepatic lipid metabolism, islet function, and energy balance(Alvarez-Dominguez et al., 2015; Chen et al., 2015; Luan et al., 2015; You et al., 2015; Zhao and Lin, 2015). Despite the fact that many studies indicate the important role of lncRNA in different tissues, little is known about the biological function of lncRNAs in chicken fat deposition, especially in chicken preadipocytes differentiation. Our study was the first to screen for lncRNAs and mRNAs regulating chicken preadipocytes differentiation by sequencing and annotating the transcriptome of four preadipocyte differentiation stages. After quality control, an average of 16.25 Gb clean reads were obtained per preadipocyte sample. A total of 1,073,884,908 reads were successfully mapped to the chicken reference genome assembly. As first study of lncRNAs in preadipocyte of chicken, we identified 27,023 novel lncRNAs. Many studies have showed that lncRNAs were fewer in exon number, shorter in length than mRNAs. Our results indicated that the predicated lncRNAs were shorter in length and contain fewer exon than mRNAs, which were in agreement with previous studies(Trapnell et al., 2010; Wang et al., 2016).

To identify genes that associated with the differentiation of preadipocytes for chicken, we compared the transcriptome-wide gene expression profiles between the libraries of the four differentiation stages. 1,336 differentially expressed lncRNAs and 1,759 differentially expressed mRNAs were obtained by pairwise comparison. To explore the stage which played crucial role in the preadipocytes differentiation, the histogram and line chart of DEGs for different stages were plotted (Figure 3E, F). The results showed that the number of differentially expressed lncRNAs and mRNAs decreased with the differentiation of preadipocytes, which suggested that the early stage might be most important for chicken preadipocytes differentiation. The differential expressed mRNAs from three comparisons (A0 vs A2, A2 vs A4, A4 vs A6) were performed to GO and KEGG analysis to explore the similarities and differences of three stages. GO analysis showed that A0-A4 stage and A4-A6 stage were similar in biological process while different from A2-A4 stage. KEGG results showed that among the three comparisons, five pathways were common, in which the JAK-STAT signaling pathway mediated the action of a variety of hormones that had profound effects on adipocyte development and function. It suggested that JAK-STAT might play important roles in the differentiation of chicken preadipocytes.

Dynamic changes of gene expression reflect an intrinsic mechanism of how an organism responds to developmental and environmental signals. Genes with a similar expression pattern are often hypothesized to have function dependence and might be coregulated by some common regulatory factors(Ye et al., 2015). Genes with known functions in a particular cluster can be a bait to point out a potential role of other function-unknown genes, facilitating the prediction of gene functions. In our study, 3,095 DEGs were successfully clustered into eight clusters. We found several interesting and important clusters associated with preadipocytes differentiation stage. Genes in K 1 cluster showed significantly and specifically up-regulated at day two of differentiation, indicating that they might greatly and specifically contribute to the starting of the preadipocytes differentiation. Genes in K6 cluster were up-regulated at day 2, 4 and 6 compared to 0 day of differentiation, suggesting their roles in the entire differentiation process. K7 cluster consisted of 307 genes that were significant up-regulated at day 6 of differentiation, demonstrating that those genes might play important roles in the later stage of differentiation. Particular attention was paid to K2 cluster which included genes that underwent an overall trend of increase, suggesting their key roles over the entire differentiation process. Members from this cluster such as *GPR39* and *CHCHD4* had been reported to regulate the differentiation of preadipocytes(Dong et al., 2016).

To investigate the function of lncRNAs, we predicted the potential targets of lncRNAs in *cis* by searching for protein-coding genes 10 kb upstream and downstream of the lncRNAs, respectively. 4,915 potential target protein-coding genes that corresponded to 10,306 lncRNAs were found and performed to GO and pathway analysis. In the biological process category, 27 terms were significantly enriched, including cellular process, system development and anatomical structure development. Besides, we found that most of the 27 significant terms were related to regulation of gene expression such as regulation of cellular process, regulation of biological process and regulation of cell communication, clearly demonstrating the role of lncRNAs in the genome. Pathway analysis showed 971 genes were enriched in 131 KEGG pathways, in which 6 pathways were associated with preadipocytes differentiation such as Wnt signaling pathway, MAPK signaling pathway and Jak-STAT signaling pathway. These results suggested that lncRNAs act on its neighoring protein-coding genesin cis to regulate abdominal preadipocytes differentiation.

Studies has shown that genes and their protein products carry out cellular processes in the context of functional modules and are related to each other through a complex network of interactions. Understanding an individual gene or protein’s network properties within such networks may prove to be as important as understanding its function in isolation(Barabasi and Oltvai, 2004; Carlson et al., 2006; Hartwell et al., 1999). Therefore, the primary emphasis in our study was on the constructing co-expression network and detecting modules related to preadipocytes differentiation. In our study, the integrated weighted gene co-expression network analysis (WGCNA) was used to construct co-expression network and detect module with 3,095 DEGs in 12 samples(Fuller et al., 2007). Using WGCNA we identified ten modules, among which six modules were stage-specific (Zhang and Horvath, 2005). It meant that those modules included genes that were down expressed or over expressed in a particular differentiation stage and can be used to represent the corresponding stage of differentiation(Jiang et al., 2014). The black, blue, green and yellow modules were correlated with stage of day two of differentiation (A2 stage), while the turquoise and brown modules were correlated with day 0 (A0 stage) and day 6 (A6 stage) of differentiation. Gene in one module suggested that they were involved in a common network of biological processes and functions. To gain insight into the biological processes and pathways involved in different stages of abdominal preadipocytes differentiation, GO and pathway analysis were performed to genes in stage-specific module. For A2 stage, given their high correlations (Figure 5C), the balck, blue, green and yellow modules were merged in GO and pathway analysis. In the GO analysis, RNA metabolic process, regulation of localization and cellular metabolic process were significantly enriched at A0 stage, generation of precursor metabolites and energy, electron transport chain and energy derivation by oxidation of organic compounds were highly enriched at A2 stage, while blastocyst development and cellular process were greatly enriched at A6 stage. Interestingly, we found that the functions of genes migrated from RNA editing at A0 stage, to generation of precursor metabolites and energy at A2 stage and finally to regulation of cell death and apoptosis at A6 stage. As far as we know, quite a few pathways involved in preadipocytes differentiation have been validated to date, including the PI3K/AKT cell signaling pathway(Dong et al., 2016), LKB1-AMPK Pathway(Tung et al., 2016), Wnt/β-catenin signaling pathway(Lu et al., 2016; Mai et al., 2014; Zhang et al., 2014c), TGF-β pathway(Park et al., 2014), Bmp/Smad pathway(Liu et al., 2014; Suenaga et al., 2013), ERK signaling pathway(Chiang et al., 2013), PDK1/Akt pathway(He et al., 2013), p38 MAPK/ATF-2 and TOR/p70 S6 kinase pathways(Yan et al., 2013). However, little is known about pathways involved in preadipocytes differentiation of chicken. In our study, many well-known pathways associated with different stages of differentiation such as ABC transporters, jak-STAT, Wnt, Insulin, Fatty acid metabolism and Fatty acid biosynthesis signaling pathways were found. Furthermore, we also identified many pathways related to preadipocytes differentiation for the first time, including Oxidative phosphorylation, Apoptosis, Propanoate metabolism and Galactose metabolism signaling pathways. The enriched KEGG pathways associated with preadipocytes differentiation in the mRNAs of stage-specific modules and potential lncRNAs targets greatly expanded our knowledge on pathways involved in preadipocytes differentiation.

To date, several previous studies have shown that PPARγ and C/EBPα(Yu et al., 2014), FATP1(Qi et al., 2013) and Klf7(Zhang et al., 2013) regulate the differentiation of preadipocytes in chicken. However, no study has been conducted on the roles of lncRNAs in chicken preadipocytes differentiation. The molecular and cellular mechanisms regulating chicken preadipocytes differentiation are still poorly understood. Here, we identified a number of highly connected lncRNAs and mRNAs in six stage-specific module. Those genes might play critical roles in each stage of preadipocytes differentiation in the chicken and were visualized by Cytoscape (Figure 7). The visualization of each module showed that the nodes number of lncRNAs were greater than that of mRNAs, demonstrating the regulation role of lncRNAs. Two lncRNAs (XLOC_068731 and XLOC_022661) and three protein-coding RNAs (*CHD6*, *LLGL1* and *NEURL1B*) were identified to be associated with day 2 of differentiation. *KLHL38* and XLOC_045161 were the highest connected mRNA and lncRNA, indicating their potential key roles in regulation of preadipocytes differentiation at day 6 of differentiation, while *ACTR6* and XLOC_070302, with the most connection, were found to be important factors in day 0 of differentiation. All the lncRNAs were novel and first identified. Those lncRNAs and mRNAs were validated by qRT-PCR and the results were in excellent agreement with the RNA-seq findings. It suggested that our findings in the present study were reliable. Besides, many well-known genes related to preadipocytes differentiation were found in the highly connected genes. For example, *IGFBP2* is a high connected gene in yellow module related to day 2 of differentiation and has inhibitory effect on preadipocyte differentiation in mice(Xi et al., 2013). It also inhibits both adipogenesis and lipogenesis in visceral(Yau et al., 2015). Similarly, *JUN* (AP-1) and *AP1B1* are highly connected genes in yellow and turquoise modules related to day 2 and day 0 of differentiation, respectively. Studies showed that *AP-1* controls adipocyte differentiation and survival by regulating *PPARγ* and hypoxia(Luther et al., 2014). *AP-1* also plays a role in regulating adipocyte differentiation and *FABP4* expression in 3T3-L1 cell(Kang et al., 2013). *AP1B1* regulates adipogenesis by mediating the sorting of sortilin in adipose tissue(Baltes et al., 2014).

## Acknowledgments

This study was financially supported by the National Broiler Industrial and Technology System: nycytx-42-G1-05 and the Priority Academic Program Development of Jiangsu Higher Education Institutions. We have received funds for covering the costs to publish in open access.

## Author Contributions

“Tao Zhang and Jinyu Wang conceived and designed the experiments; Tao Zhang and Xiangqian Zhang performed the experiments; Tao Zhang and Genxi Zhang analyzed the data; Kaizhou Xie, Kenpeng Han and Qian Xue contributed reagents/materials/analysis tools; Tao Zhang and Jinyu Wang wrote the paper.”

## Abbreviations

lncRNA: Long Non-coding RNA
DEGs: Differentially Expressed Genes
IGFBP2: Insulin like growth factor binding protein 2
JUN: Jun proto-oncogene
CDS: Coding sequence
UTR: Untranslated Regions
GO: Gene Ontology
KEGG: Kyoto Encyclopedia Of Genes And Genomes
WGCNA: Weighted Gene Co-expression Network Analysis
FDR: False Discovery Rate
GPR39: G protein-coupled receptor 39
AP-1: Transcription factor subunit
CHCHD4: Coiled-coil-helix-coiled-coil-helix domain containing 4
AP1B1: Adaptor related protein complex 1 beta 1 subunit
FABP4: Fatty acid binding protein 4
FBS: Fetal bovine serum
IBMX: 3 isobutyl 1 methylxanthine

